# Cerebral Organoids In Primary Progressive Multiple Sclerosis Reveal Stem Cell Disruption And Failure To Produce Oligodendrocytes

**DOI:** 10.1101/2022.06.24.497517

**Authors:** Nicolas Daviaud, Eric Chen, Tara Edwards, Saud A Sadiq

## Abstract

Multiple sclerosis (MS) is an auto-immune inflammatory disorder affecting the central nervous system. The cause of the disease is unknown but both genetic and environmental factors are implicated in the pathogenesis. We derived cerebral organoids from induced pluripotent stem cells (iPSC) of healthy control subjects as well as from primary progressive MS (PPMS), secondary progressive MS (SPMS) and relapsing remitting MS (RRMS) patients to better understand the pathologic basis of the varied clinical phenotypic expressions of MS. In MS organoids, most notably in PPMS, we observed a decrease of proliferation marker Ki67 and a reduction of the SOX2^+^ stem cell pool associated with an increased expression of neuronal markers CTIP2 and TBR1. This dysregulation of the stem cell pool is associated with a decreased expression of the cell cycle inhibitor p21. Our findings show that the genetic background of a patient can directly alter stem cell function. This study also provides new insights on the innate cellular dysregulation in MS and identifies p21 pathway as a new potential target for therapeutic strategies in MS.

**Summary Statement:** Using cerebral organoids derived from patients with multiple sclerosis we detected that p21 decrease may induce a disruption of the stem cell cycle leading to a defect of oligodendrocyte differentiation

## INTRODUCTION

Multiple sclerosis (MS) is an auto-immune inflammatory disorder associated pathologically with widespread focal lesions of primary demyelination with variable axonal, neuronal and astroglia injury in the brain and the spinal cord. Clinically, it may progress to irreversible neurological disability and cognitive decline (Filippi et al., 2018).

The major clinical subtypes of MS at initial presentation are RRMS and PPMS. RRMS represents the initial inflammatory phase of approximately 85–90% of all cases. Over time this form may evolve to SPMS usually with progressive failure of remyelination and with associated axon degeneration (Lublin and Reingold, 1996). PPMS affects approximately 10% of all cases and manifests with a steady decline in function that occurs from disease and symptom onset (Andersson et al., 1999) notably with an absence of relapses and remissions. The origin and evolution of MS is still poorly understood partly due to the relative inaccessibility of human brain tissues and inadequate animal models to study the disease (Ransohoff, 2012). Furthermore, the physiological basis of the varied phenotypic expression of MS subtypes is unknown.

It is now accepted that MS development is influenced by both genetic factors. Familial relatives of patients with MS, especially first-degree relatives, are more susceptible to develop MS compared to the general population. Monozygotic twins have higher risks of getting MS (up to 30%), non-twin siblings have a risk around 1%, while the general population has only 0.1% of chance of getting MS (Sadovnick et al., 1998; Willer et al., 2003; O’Gorman et al., 2013; Westerlind et al., 2014). Moreover, risk variants influenced the expression of 203 genes and 21% of intergenic variants are associated with gene regulation in cortex tissue but not in immune cells (International Multiple Sclerosis Genetics, 2019).

Recent advances in 3D cerebral organoid (c-organoids) cultures derived from induced pluripotent stem cells (iPSC) provide new avenues to implement reproducible models to study cell type- and stage-specific effects of MS. C-organoids contain ventricle-like structured aligned by neural stem cells, progenitor cells in various stages of differentiation and migration, glial cells and cortical neurons in a stereotypical inside-out stratified layout (Lancaster and Knoblich, 2014; Daviaud et al., 2018; Daviaud et al., 2019). Moreover, neurons present in c-organoids are able to get myelinated (Madhavan et al., 2018; Matsui et al., 2018). We sought to develop an innovative model of MS using human iPS derived c-organoids. C-organoids developed from MS patient induced-pluripotent stem cells (iPSCs) provides insight into the effect of patient genetic background on neural cells and their interactions in a model free of immune system interaction and within a controlled microenvironment.

We report as a proof of concept, the derivation of c-organoids from iPSCs of healthy control subjects as well as from PPMS, SPMS and RRMS patients. Using this innovative c-organoid model, a dysregulation of the NPC proliferation and differentiation capacity, resulting in a decrease of proliferation marker Ki67, as well as a reduction of the SOX2^+^ stem cell pool is observed, most noticeably in PPMS. This decrease of proliferation and stemness is associated with a larger cortical plate and an increased expression of neuronal markers CTIP2 and TBR1 in MS organoids. This proliferation/differentiation imbalance might be due to a shift of cell division mode for neurogenesis. In addition, a strong decrease of the cell cycle inhibitor p21 expression in the ventricular zone (VZ) is observed in PPMS, in a p53/apoptosis independent pathway, thus affecting directly the cell cycle and the cell division mode.

Our studies with c-organoids in MS provide novel insights into the development of neural interactions within the defined genetic background of MS patients. This study suggests that there is an innate cellular dysregulation in MS and implicate dysfunction of the p21 pathway as a critical abnormality in PPMS.

## RESULTS

### Establishment and characterization of blood cell derived IPS cells and cerebral organoids

To develop an experimental model of MS focusing on patient genetic background, the protocol developed by Lancaster and colleagues was used (Lancaster et al., 2013; Daviaud et al., 2019), with minor modifications, to derive consistent c-organoids from human iPSCs with dorsal forebrain specification (Fig. S2A). Over the next 4 weeks of culture, we observed rapid maturation of c-organoids with appearance of ventricle like structures aligned with a VZ-like proliferative zone containing Ki67^+^ proliferative cells and SOX2^+^ neural precursors pool (Fig. S2B and S2C) and a rudimental CP containing DCX^+^ neuroblast, TBR1^+^ & CTIP2^+^ neurons (Fig. S2B and S2C), separated by an SVZ-like transitional zone containing TBR2^+^ (EOMES) intermediate progenitor cells (IPC) (Fig. S2B, S2C).

### Cerebral organoid growth rate assessment

C-organoids were observed under inverted microscope regularly during the 42 days of culture to identify any difference in growth rates, particularly during the early stages of development (Fig. 1A). Organoids measured an average of 250 µm diameter on the 1^st^ day of culture and reached about 450 µm at D4 and over 1 mm diameter at D14. Organoid surface was analyzed at D2 and D9 (Fig. 1B). After 2D of culture in vitro, control organoids measured 0.16 mm² while PPMS organoids were significantly bigger, reaching 0.21 mm² (Kruskal-Wallis test, p = 0.0012). RRMS organoids and SPMS organoids were not significantly different than controls with 0.18 mm² (Kruskal-Wallis test, p = 0.5710) and 0.14 mm² (Kruskal-Wallis test, p = 0.7294) respectively. After 9 days of culture all organoids reached a similar size around 0.65 mm² with no significant difference detected between the different conditions (Fig. 1B).

**Figure 1:**
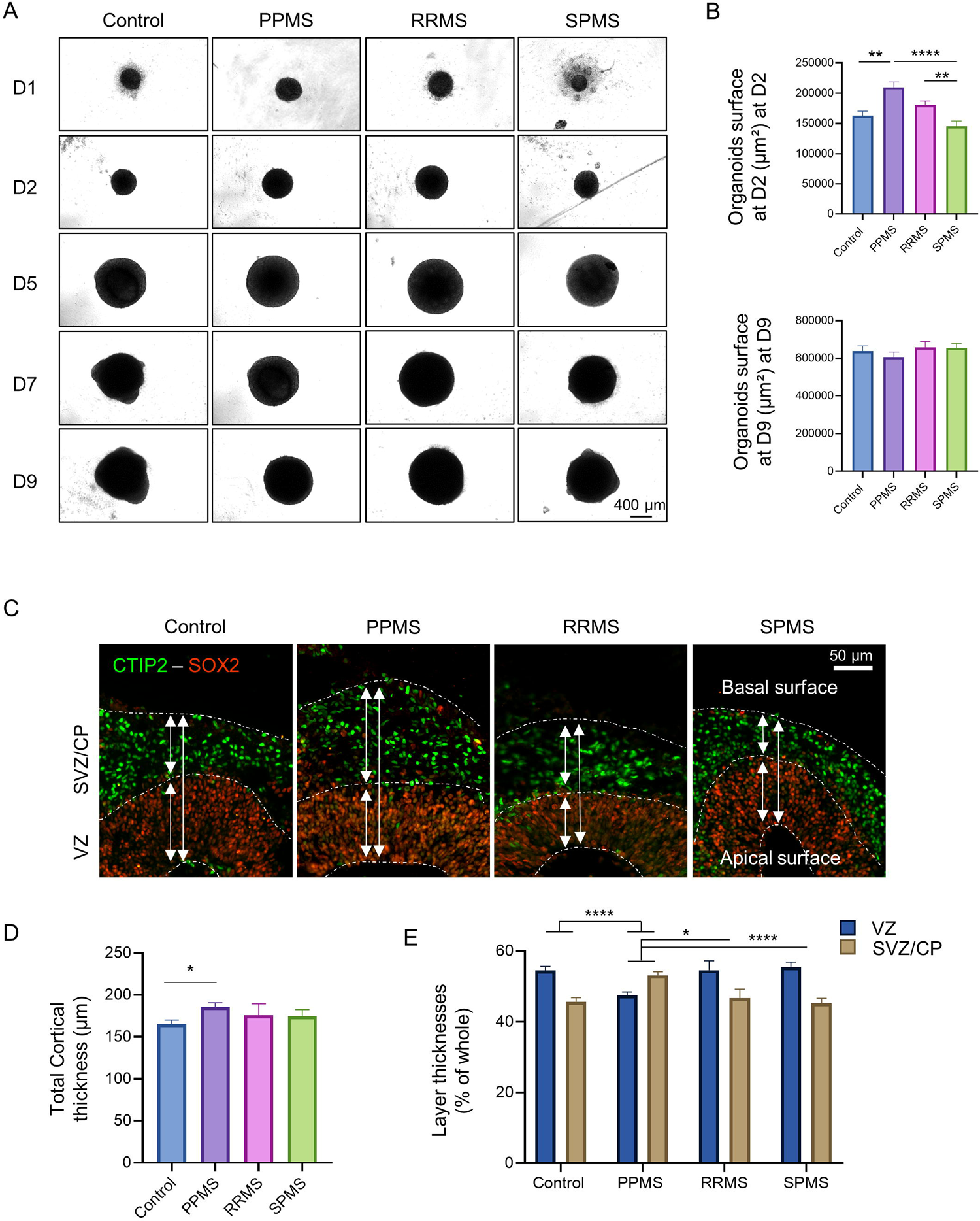
MS organoids display altered growth rate in culture. A) Brightfield pictures of c-organoids derived from control and MS patients in culture from day 1 to day 9. B) Measurements of c-organoids surface in µm² at D2 (top graph) and at D9 (bottom graph). Analysis showed significant larger PPMS organoids compared to control at D2 only. Kruskal-Wallis test (H value = 31.30, p < 0.0001) followed by a Dunn’s post hoc test. No significant difference was detected at D9. Kruskal-Wallis test (H value = 2.88, p = 0.4098). C) Immunofluorescence of c-organoids at D42 for the stem cell marker SOX2 and neuronal marker CTIP2. The staining shows a neat separation between the VZ and the SVZ/CP in the c-organoids cortical structure. D) Measurement of the c-organoids cortical structure thickness for control and MS subtypes at D42. Quantification revealed a significantly thicker cortical structure in PPMS compared to control. Kruskal-Wallis test (H value = 8.323, p = 0.0398) followed by a Dunn’s post hoc test. E) Measurement of the thickness of the 2 main layers of the cortical structure, the VZ and the SVZ/CP at D42. Quantification showed that PPMS organoids possessed a significantly larger SVZ/CP and a thinner VZ compared to other conditions. Two-way ANOVA (F (3, 532) = 20.13, p <0.0001) followed by a Tukey post hoc test. *: p < 0.05, **: p < 0.01, ***: p < 0.001, ****: p < 0.0001

After 42 days in culture, c-organoids were sliced and analyzed by immunofluorescence to determine the thickness of the cortical structure (Fig. 1C). Control cortical structures were +/- 165 µm thick. PPMS cortical structures were significantly thicker reaching 185 µm (Kruskal-Wallis test, p = 0.026) while RRMS and SPMS cortical structure reached 175 µm (p > 0.99) and 174 µm (Kruskal-Wallis test, p > 0.99) respectively but the difference was not significant (Fig. 1D).

In the cortical structure, the two main cortical layers, the VZ and the SVZ/CP can be easily identified by SOX2 and CTIP2 staining respectively (Fig. 1C). In control, RRMS and SPMS organoids, the VZ represented an average of 55-54% of the cortical structure while the SVZ/CP represented 45-46%. PPMS organoids were significantly different from control, RRMS and SPMS, as the VZ represented +/- 47% of the cortical structure (Two-way ANOVA, p = 0.0120 vs control) while the CP represented 53% of the cortical structure (Two-way ANOVA, p < 0.0001 vs control) (Fig. 1E). This result might indicate a dysregulation of the stem cell proliferation/differentiation capacity in PPMS organoids.

### MS organoids display a disruption of the stem cell proliferation capacity

To assess the effect of MS genetic background on progenitor cell proliferation, immunofluorescence against proliferation marker Ki67 was performed in c-organoids at D42. As expected, proliferating Ki67^+^ cells were localized alongside the ventricle apical surface (Fig. 2A) and no ectopic location of Ki67^+^ cells were detected. A quantification of the number of Ki67^+^ cells showed a significant decrease of Ki67^+^ cell number in PPMS compared to control (Kruskal-Wallis test, p = 0.0309) and compared to RRMS (Kruskal-Wallis test, p = 0.0245) while no difference was observed with SPMS (Kruskal-Wallis test, p = 0.211) (Fig. 2B).

**Figure 2:**
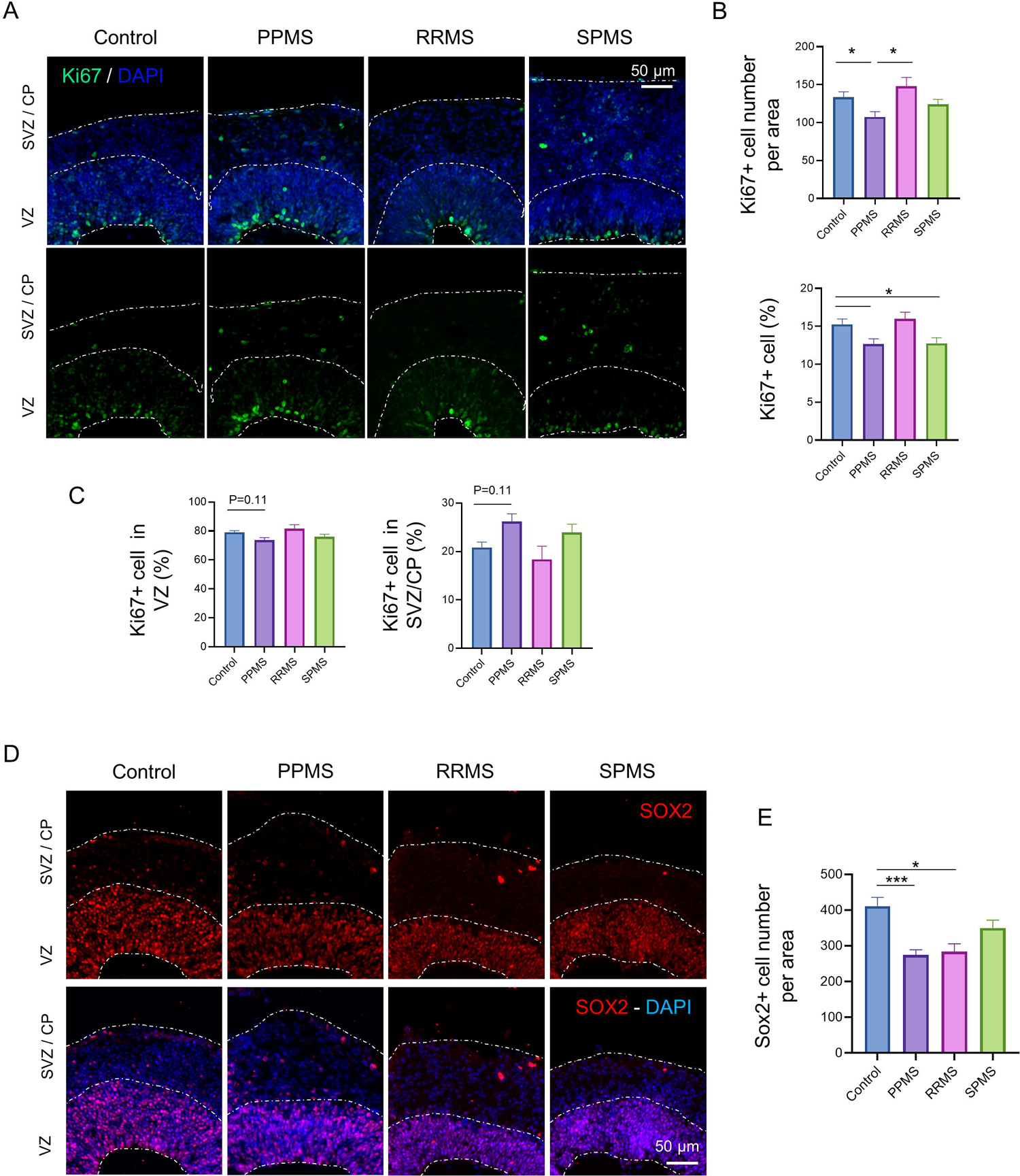
Impairment of stem cell population in MS organoids. A) Representative images of an immunofluorescence against the proliferation marker Ki67 in c-organoids at D42. B) Quantifications of Ki67^+^ cells number and percentage of Ki67^+^ cells in the organoids cortical structure at D42. Quantifications revealed a decrease of Ki67^+^ number (Kruskal-Wallis test, H value = 11.77, p = 0.0082, followed by a Dunn’s post hoc test) as well as percentage in PPMS compared to control (One-way ANOVA, F (3, 217) = 4.274, p = 0.0059, followed by a Tukey’s multiple comparisons test). C) A quantification of the percentage of Ki67^+^ cells in the VZ (radial glia) and the SVZ/CP (intermediate progenitor cells and neuroblast) was performed. A slight decrease was detected in the VZ while an increase was observed in the SVZ/CP (Kruskal-Wallis test, H value = 8.38, p = 0.0387 followed by a Dunn’s post hoc test). D) Representative images of an immunofluorescence against the NPC marker SOX2 in c-organoids at D42. E) Quantification of SOX2^+^ cells in c-organoids cortical layers showed a significant lower SOX2^+^ cell number in PPMS and RRMS compared to control organoids (One-way ANOVA, F (3, 138) = 6.191, p = 0.0006, followed by a Tukey post hoc test). To ensure quantification consistency, analyzed cortical area were cropped from the original picture to a 250 µm wide x 300 µm image, spanning all cortical layers (VZ, SVZ, and CP). *: p < 0.05, **: p < 0.01, ***: p < 0.001, ****: p < 0.0001

To make sure the decrease of Ki67^+^ cells observed was not due to a reduction of the size of the VZ, a quantification of the percentage of Ki67^+^ per DAPI was also performed. A lower percentage of Ki67^+^ cells was measured in PPMS (One-way ANOVA, p = 0.0410) and SPMS (One-way ANOVA, p = 0.0438) compared to control while no difference was observed with RRMS (One-way ANOVA, p = 0.9588), confirming our previous result (Fig. 2B).

Cortical c-organoids hold two main population of dividing cells, the NPC localized at the apical surface of the VZ and the IPCs localized in the SVZ. Quantification of each population of proliferating cell was assessed. A slight decrease of Ki67 proliferation marker was detected in the VZ (Kruskal-Wallis test, p = 0.11) while a slight increase was measured in the SVZ/CP (Kruskal-Wallis test, p = 0.11) in PPMS organoids compared to control, while no difference was observed with the other MS subtypes (Fig. 2C).

These results indicate that a decrease of proliferation occurs in PPMS compared to control. This reduction is noted particularly with the NPCs, while an increase proliferation is observed with IPCs. This might indicate a reduction of stem cell proliferation capacity and an increase of neurogenesis.

### Stem cell pool is reduced in MS organoids

As a reduction of the number of proliferative cells was detected in MS organoids, an immunofluorescence was performed against stem cell marker SOX2 to assess the stem cell pool status. As expected, majority of SOX2^+^ cells were present in the VZ, while a few cells were positive in the upper layers (Fig. 2D). No difference of SOX2^+^ cell location was detected however, quantification showed a significant decrease of SOX2^+^ cell number in the cortical structure most notably of PPMS organoids (One-way ANOVA, p = 0.0006) and less so in RRMS (One-way ANOVA, p = 0.0363) compared to control, while no significant difference was detected with SPMS (One-way ANOVA, p = 0.3217) (Fig. 2E). This result indicates that there is a decreased stem cell pool associated with a decrease of proliferation in MS organoids, most evidently seen in PPMS.

### Neural progenitors are increased in MS organoids

To observe the effects of cell cycle dysregulation on differentiation, immunofluorescence was performed on two population of precursors, TBR2^+^ intermediate progenitors and DCX^+^ neuroblasts. TBR2 expression marks the transition from radial glia to intermediate progenitors (Englund et al., 2005; Sessa et al., 2008), therefore it was assumed that a reduction of SOX2^+^ radial glia might contribute to a decrease of TBR2^+^ IPCs. TBR2 staining was observed along the SVZ, with no obvious alterations between the different conditions (Fig. 3A). Quantification showed a slight decrease of TBR2^+^ cell number in PPMS and RRMS organoids compared to control but didn’t reach significance (One-way ANOVA, p = 0.19 and p = 0.35 respectively) while no difference was observed with SPMS organoids (One-way ANOVA, p = 0.99) (Fig. 3B).

**Figure 3:**
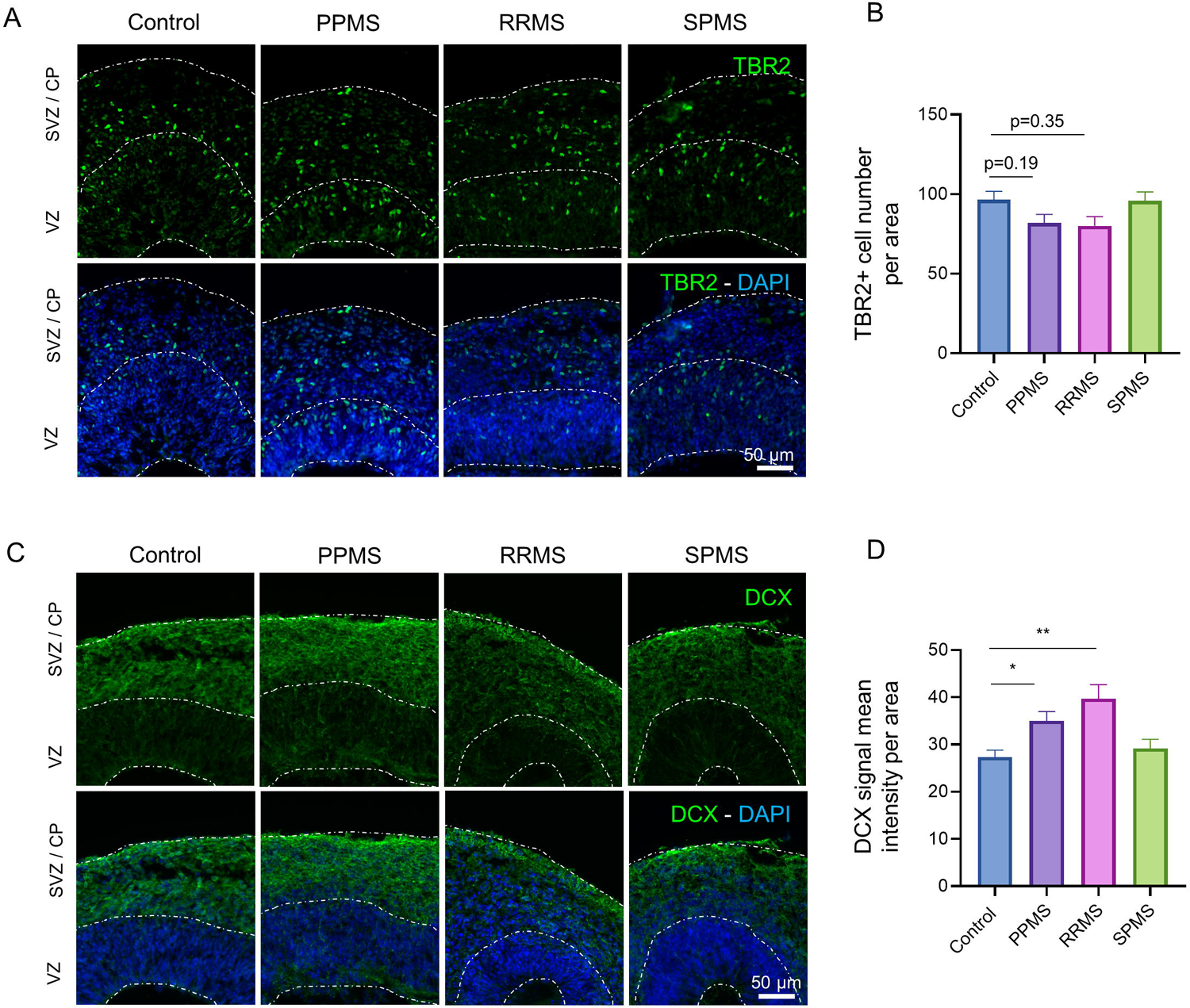
MS organoids exhibits larger neural progenitor population. A) Representative images of an immunofluorescence against the IPC marker TBR2 in c-organoids at D42. B) Quantifications of TBR2^+^ cell number in cortical structure of organoids at D42 did not show any significant difference between the different conditions (One-way ANOVA, F (3, 149) = 2.162, p= 0.0949, followed by a Tukey post hoc test). C) Representative images of an immunofluorescence against the neuroblast marker DCX in c-organoids at D42. D) DCX immunofluorescence mean intensity was measured and compared in each condition. A significant increase of DCX immunostaining intensity was measured in PPMS and RRMS compared to control, while no difference was observed for SPMS organoids (Kruskal-Wallis test, H value = 18.01, p = 0.0004, followed by a Dunn’s post hoc test). To ensure quantification consistency, analyzed cortical area were cropped from the original picture to a 250 µm wide x 300 µm image, spanning all cortical layers (VZ, SVZ, and CP). *: p < 0.05, **: p < 0.01, ***: p < 0.001, ****: p < 0.0001

Doublecortin (DCX) is a microtubule-associated phosphoprotein that promotes neurite extension and cell migration in the cortex (Gleeson et al., 1999). As expected, DCX staining showed that DCX^+^ cells were localized only in the SVZ/CP of the cortical organoids (Fig. 3C). DCX immunofluorescence mean intensity was measured with a significant increase of DCX immunofluorescence intensity seen in PPMS (Kruskal Wallis, p = 0.0156) and RRMS (Kruskal Wallis, p = 0.0015) compared to control. No significant difference in intensity was observed in SPMS organoids (Kruskal Wallis, p > 0.9999) compared to control (Fig. 3D).

### MS organoids display an increased neuronal differentiation and decreased oligodendroglia cell population

To assess the cortical neuron population in MS organoids compared to control, immunostainings for CTIP2 and TBR1 were performed. CTIP2 is a marker expressed in excitatory neurons localized in the deep neocortical layer (Nadadhur et al., 2017) while TBR1 is particularly expressed in glutamatergic pyramidal neurons in the cerebral cortex (Hodge et al., 2012). As expected, CTIP2 and TBR1 were expressed mostly in the outer layers of c-organoids (Fig. 4A and 4C), no ectopic localization of neuronal markers was detected in the different conditions.

**Figure 4:**
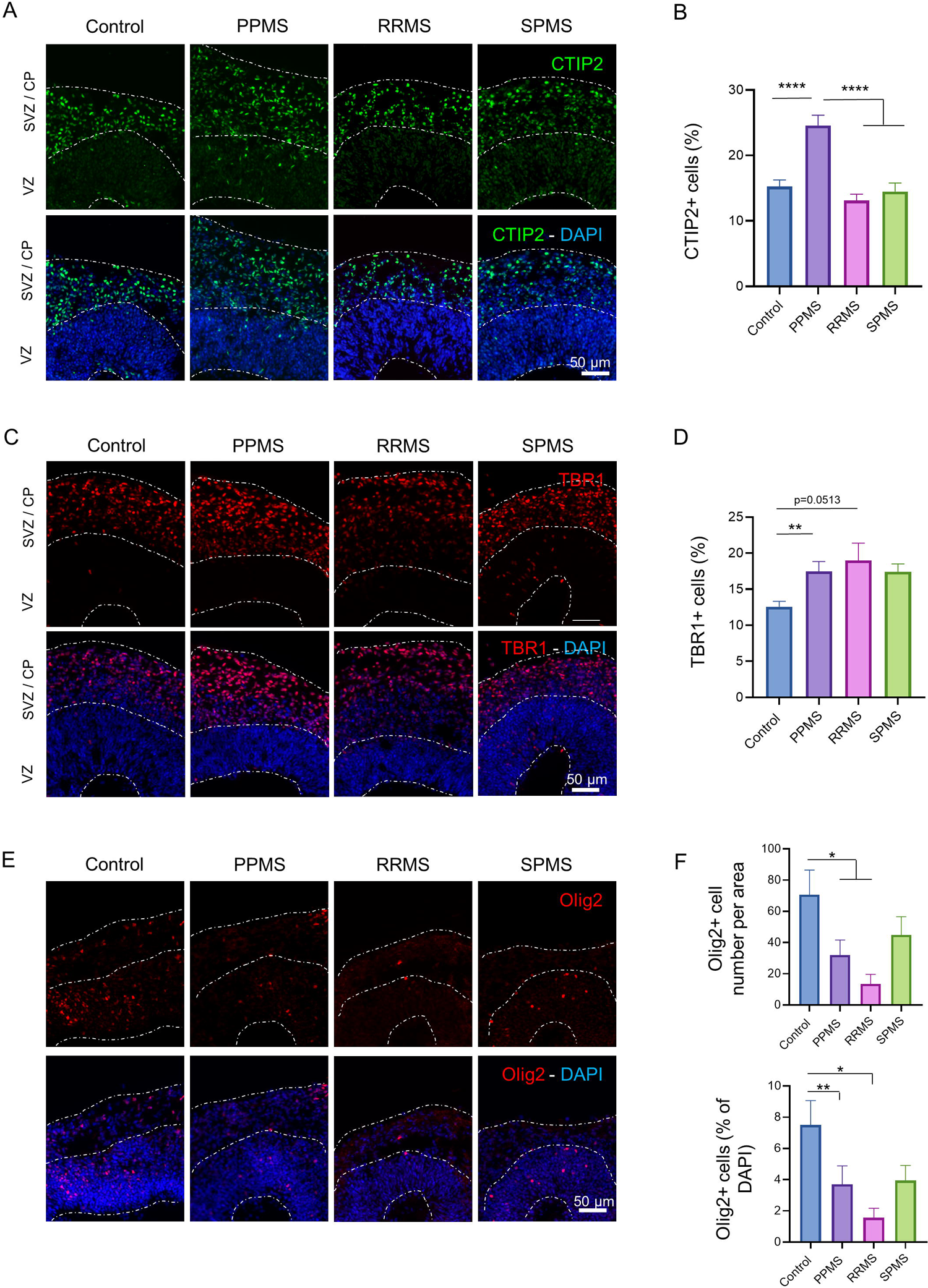
MS organoids display an increased number of mature neurons and a reduced oligodendrocyte population. A) Representative images of an immunofluorescence against the deep-layer cortical neuron marker CTIP2 in c-organoids at D42. B) Quantification of the percentage of CTIP2^+^ cells in c-organoids revealed a significant higher expression of CTIP2 in PPMS compared to control, RRMS and SPMS (Kruskal-Wallis test, H value = 36.94, p < 0.0001, followed by a Dunn’s post hoc test). C) Representative images of an immunofluorescence against the early-born cortical neuron marker TBR1 in c-organoids at D42. D) Quantification of TBR1^+^ cell percentage in organoid cortical structure showed a significant higher expression of TBR1^+^ cells in PPMS compared to control (One-way ANOVA, F (3, 94) = 4.701, p= 0.0042, followed by a Tukey’s multiple comparisons test). E) Immunofluorescence pictures of oligodendrocyte precursors (Olig2) in c-organoids at D42. Olig2^+^ cells were mostly localized in the VZ. F) Quantification of Olig2^+^ cell number per area and of the percentage of Olig2^+^ cells per area. Quantification revealed a significant lower number of Olig2^+^ cells in PPMS and RRMS organoids compared to control (Kruskal-Wallis test, H value = 14.07, p < 0.0028, followed by a Dunn’s post hoc test), as well as a lower percentage of Olig2^+^ cells in PPMS and RRMS compared to control (Kruskal-Wallis test, H value = 14.77, p < 0.0020, followed by a Dunn’s post hoc test). To ensure quantification consistency, analyzed cortical area were cropped from the original picture to a 250 µm wide x 300 µm image, spanning all cortical layers (VZ, SVZ, and CP). *: p < 0.05, **: p < 0.01, ***: p < 0.001, ****: p < 0.0001

A significant higher percentage of CTIP2^+^ cell per cortical area was detected in PPMS compared to control (Kruskal Wallis, p < 0.0001), RRMS (Kruskal Wallis, p < 0.0001) and SPMS (Kruskal Wallis, p < 0.0001) (Fig. 4B). Analysis of TBR1 expression showed a significant higher percentage of TBR1^+^ cells in PPMS compared to control (One-way ANOVA, p = 0.0093). No difference with the other MS subtypes was detected (One-way ANOVA, p = 0.0513 and p = 0.1153 for RRMS and SPMS respectively) (Fig. 4D). This result might indicate that the detected increase of neurogenesis might particularly affect excitatory neurons localized of the neocortical layer.

An immunofluorescence for GABAergic neuronal marker GAD67 and glutamatergic neuronal marker vGluT1 was performed on c-organoids. Both neuronal markers could be found in organoids derived from healthy and MS patients (Fig. S3) with no obvious expression difference.

Oligodendrocytes are responsible for myelin sheet formation during brain development. To study oligodendroglia cell population an immunofluorescence was performed for the oligodendrocyte transcription factor 2 (Olig2), an oligodendrocyte marker. Olig2 was localized in the VZ of organoid cortical structure (Fig. 4E), no ectopic location of Olig2^+^ cells were detected in the different MS subtypes. Quantification revealed a significant lower number of Olig2^+^ cells in PPMS organoids (Kruskal Wallis, p = 0.0409) as well as RRMS organoids (Kruskal Wallis, p = 0.0383) compared to control. To further confirm this result, the percentage of Olig2^+^ cells was measured as well. A significant lower percentage of Olig2^+^ cells was measured in PPMS (Kruskal Wallis, p = 0.0063) and RRMS compared (Kruskal Wallis, p = 0.0499) to control (Fig. 4F). This suggests that remyelination capacity is innately diminished in PPMS patients.

### Cell cycle inhibitor p21 expression is drastically reduced in PPMS organoids

To understand the mechanisms associated with the disruption of the stem cell pool, an analysis of p21 expression was performed. p21 is a cyclin-dependent kinase inhibitor that plays essential roles in the cell cycle regulation, proliferation, differentiation as well as DNA damage response and apoptosis (Cazzalini et al., 2010; Kreis et al., 2019) and can specifically binds to SOX2 (Marques-Torrejon et al., 2013). Immunofluorescence in c-organoids revealed that p21 was mostly expressed in the VZ with a few cells localized in the SVZ/CP. However, almost no p21-positive cells were found in PPMS c-organoids (Fig. 5A). Quantification showed that an average of +/- 16% of cortical cells expressed p21 in control organoids. A dramatic decrease of p21^+^ cells was detected in PPMS, reaching about 4.5% (Kruskal Wallis, p < 0.0001) compared to control. SPMS also exhibited a significant lower percentage of p21^+^ cells compared to control organoids, reaching +/- 10% (Kruskal Wallis, p < 0.0001) while RRMS exhibited only a slight decrease, reaching +/- 11% (Kruskal Wallis, p = 0.1269). It is interesting to note that PPMS also displayed a significant lower percentage of p21^+^ cells than RRMS (Kruskal Wallis, p = 0.0006) and SPMS (Kruskal Wallis, p < 0.0001) (Fig. 5B).

**Figure 5:**
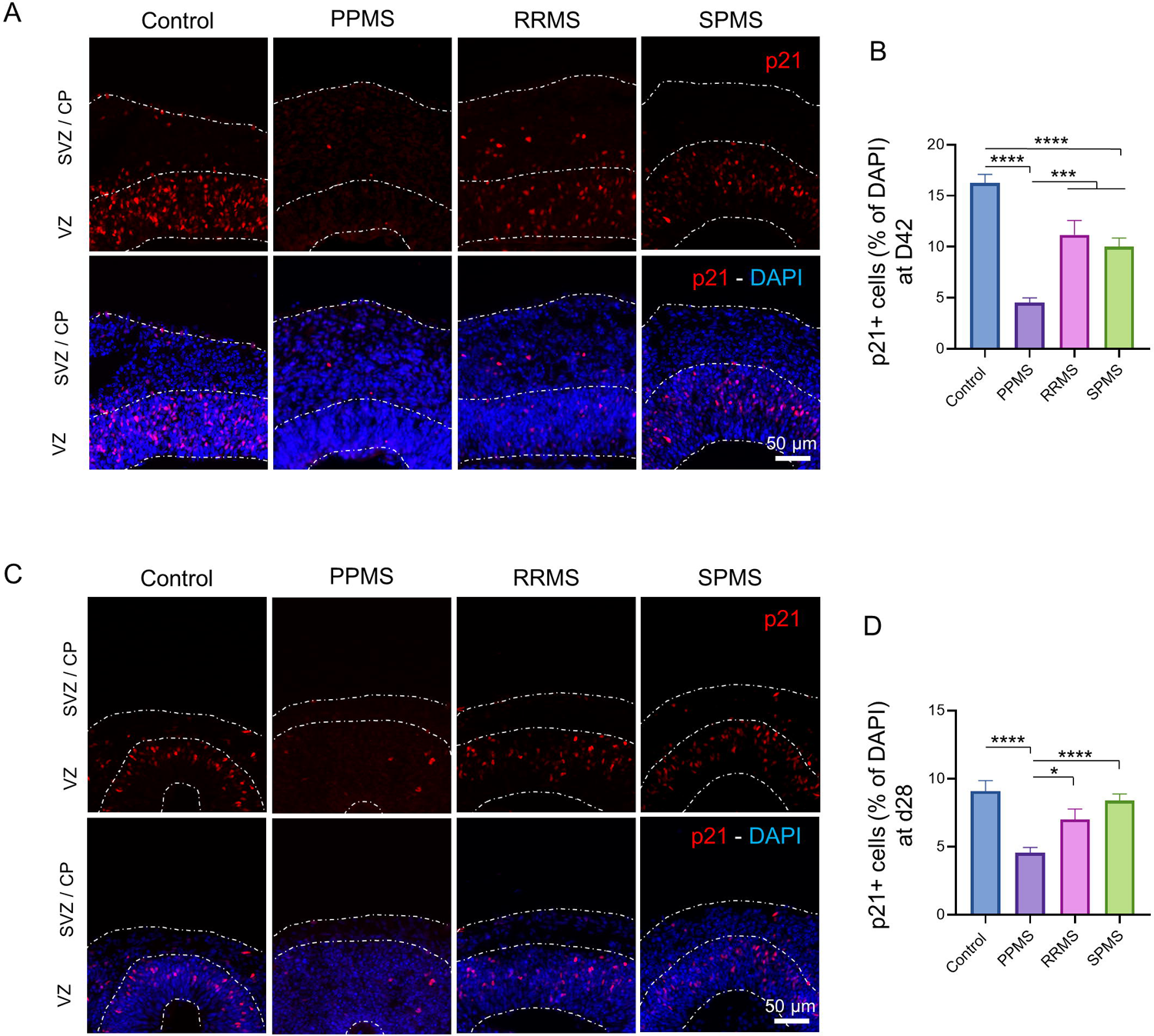
p21 expression is drastically reduced in PPMS organoids. A) Representative images of an immunofluorescence against the cyclin-dependent kinase inhibitor marker p21 in c-organoids at D42. p21^+^ cells were mostly expressed in the VZ and the SVZ. B) p21^+^ cells percentage was counted in the c-organoids cortical structure. A strong and significant decrease of p21 expression was detected in PPMS compared to control, RRMS and SPMS (Kruskal-Wallis test, H value = 95.64, p < 0.0001, followed by a Dunn’s post hoc test). C) Representative images of an immunofluorescence against p21 in c-organoids at D28. p21^+^ cells were mostly expressed in the VZ. D) Quantification of p21^+^ cells in c-organoid cortical structures. A significant decrease of p21 percentage was detected in PPMS compared to control, RRMS and PPMS. (Kruskal-Wallis test, H value = 53.02, p < 0.0001, followed by a Dunn’s post hoc test). To ensure quantification consistency, analyzed cortical area were cropped from the original picture to a 250 µm wide x 300 µm image, spanning all cortical layers (VZ, SVZ, and CP). *: p < 0.05, **: p < 0.01, ***: p < 0.001, ****: p < 0.0001

p21 expression analysis was also performed in organoids at D28, instead of D42, to be able to verify if this reduction of p21 occurred already in early time-point. At day 28, the cortical plate is not very developed yet, and organoid cortical structures are mostly composed of stem/precursor cells. p21 was detected in the VZ of control organoids but was decreased in PPMS organoids (Fig. 5C). Quantification revealed a significant reduction of percentage of cell expressing p21 in PPMS organoids compared to control (Kruskal Wallis, p < 0.0001), while no significant difference was observed with RRMS (Kruskal Wallis, p = 0.8588) and SPMS (Kruskal Wallis, p > 0.9999) compared to control (Fig. 5D). Interestingly PPMS organoids also contained significantly less p21^+^ cells than RRMS and SPMS (Kruskal Wallis, p = 0.0124 and p < 0.0001 respectively). This result suggest that the disruption of the cell cycle and reduction of proliferative capacity might be mediated by p21 loss in MS c-organoids.

### DNA damage and apoptosis pathway are not involved in organoid cell population fate in PPMS

Expression of p21 can be up-regulated by the p53 tumor suppressor gene in vitro in response to DNA-damaging agents. However, p21 expression can be regulated independently of p53 in normal tissue during cell growth, differentiation, and also following DNA damage (Macleod et al., 1995) (Fig. 6A). To assess whether or not p21 expression is followed by DNA damage and associated with the apoptosis pathway, an immunofluorescence was performed against DNA damage marker γH2AX, tumor suppressor gene P53 and apoptosis activator cleaved caspase 3 (CC3) in control and PPMS organoids at D42 (Fig. 6B).

**Figure 6:**
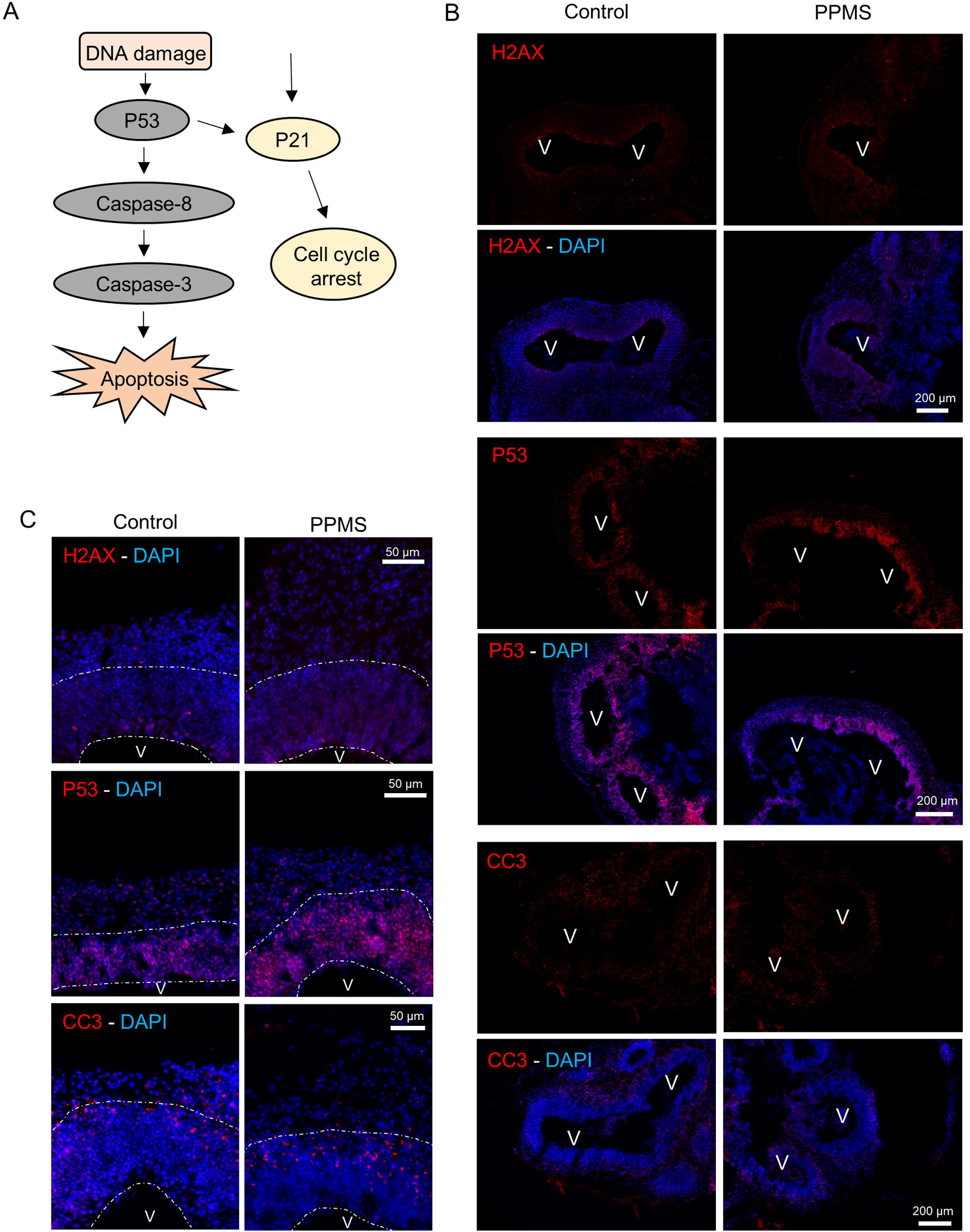
DNA damage pathway involvement in MS organoids. A) Schematic description of the DNA damage pathway, induced by DNA damage and leading to apoptosis. B) Immunofluorescences of c-organoids at D42 for DNA damage marker γ p53 and apoptosis marker cleaved caspase 3 (CC3) in PPMS and healthy controls. V: ventricle C) Immunofluorescence for γH2AX, p53 and CC3 in control and PPMS organoids at D42 in cropped images of 250 µm wide x 300 µm, spanning all cortical layers (VZ, SVZ, and CP). No noticeable difference was observed in the different marker between control and MS cortical structure. V: ventricle

H2AX was expressed in a low number of cells in control organoids as well as PPMS organoids. The majority of the H2AX^+^ cells were localized on the apical surface of the ventricle. No dramatic difference was observed between the two conditions. (Fig. 6B and C). P53 immunofluorescence revealed that a high number of p53^+^ cells were expressed in control and PPMS organoids. P53^+^ cells were mostly localized in the VZ, where the stem cell pool is also located, with little positive cell localized in the SVZ/CP. The amount of p53^+^ cells seemed similar in PPMS and control organoids. (Fig. 6B and C). CC3 shows cells undergoing apoptosis. A low number of CC3^+^ cells were observed in control and PPMS organoids. CC3^+^ cells were mostly localized in the SVZ/CP, indicating that a few neurons are undergoing apoptosis, while a few to no positive cells were detected in the VZ. No difference was observed between control and PPMS organoids. (Fig. 6B and C).

Overall, these results indicate that the change in p21 expression and modification of the cell cycle observed in PPMS is likely independent of the DNA damage/apoptosis pathway and due to an inner genetic or epigenetic variation.

### Cleavage plane angle shifts in MS organoids

Neural stem cells can expand through symmetric divisions with vertically oriented cleavage planes relative to the apical ventricular surface or undergo neurogenesis through asymmetric divisions with horizontally or oblique oriented cleavage planes (Fig. 7A) (Chenn and McConnell, 1995; Mione et al., 1997). To assess whether HI affects cell division mode, NPC were stained for G2/M phase marker phospho-histone H3 (PH3) in c-organoids from all the different subtypes of MS as well as control (Fig. 7B).

**Figure 7:**
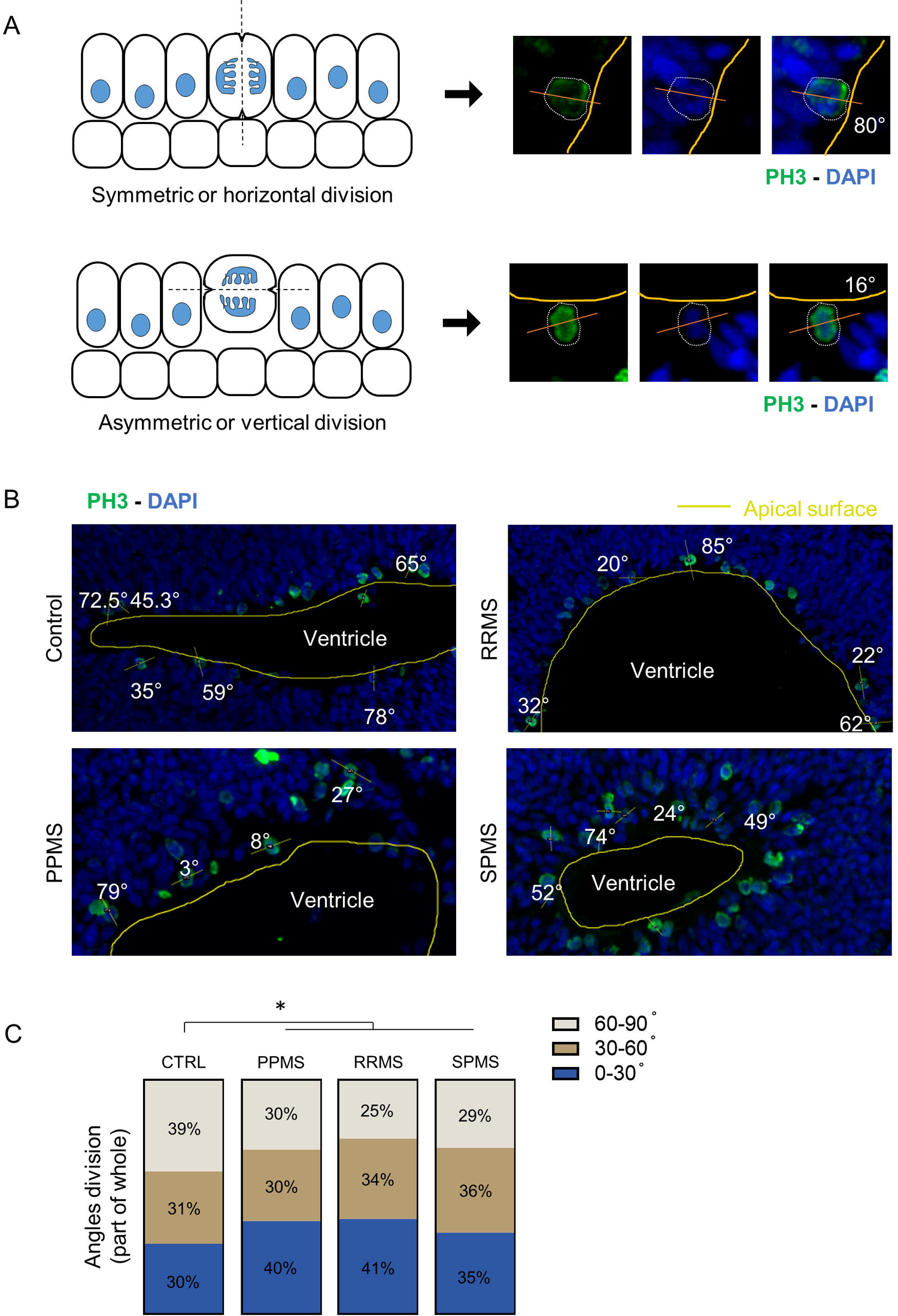
Radial glia division mode shift to asymmetric in PPMS. A) Schematic representation of symmetric (horizontal) and asymmetric (vertical) division modes (left). Illustrative pictures of mitotic cells, identified by PH3 immunofluorescence, and their division angle measurements. Yellow line represents the apical surface of the ventricle, white dotted line represents the cell edge, the orange line represents the cell division axis. B) Representative pictures of PH3 immunofluorescence in c-organoids at D42 in the different conditions. Measurement of division angle of mitotic cell was measured against the apical surface of the ventricle. Yellow lines represent the ventricle apical surface, orange lines represent the mitotic cell division axis. Angle are shown in degree. C) Representation of the different angle division modes in part of whole measured in c-organoids at D42. 0-30° division mode represents a symmetric division (stemness) while 30-60° and 60-90° represent asymmetric division (neurogenic). A significant increase of asymmetric division was detected in all MS conditions compared to control. Chi-Square test; About 20 cells were count per batch, 4 different batches containing 3-4 organoids were analyzed. *: p < 0.05, **: p < 0.01, ***: p < 0.001, ****: p < 0.0001

The incidence of horizontal (0-30°), oblique (30-60°) and vertical division (60-90°) was measured. In control, an average of 40% of cell were dividing following a horizontal cleavage while this percentage dropped to +/- 29.7% in PPMS samples (Chi-square test, p = 0.0374), 25% in RRMS (Chi-square test, p < 0.0001) and 28.9% in SPMS (Chi-square test, p < 0.0001) (Fig. 7C). This result shows a change of the angle of division in MS organoids compared to control, which indicates a reduction of NPC proliferative capacity, in favor of neurogenesis, which may result in a reduction of the progenitor pool and an expansion of neuronal populations.

### Cultured IPS derived neural progenitors exhibit senescence and premature differentiation

To further investigate the effect of genetic background of patient with MS on neural progenitors, patient iPSCs were derived into neural progenitor cells in vitro and analyzed by immunofluorescence during expansion and after 10 days of differentiation by mitogen withdrawal (Fig. 8A-B).

**Figure 8:**
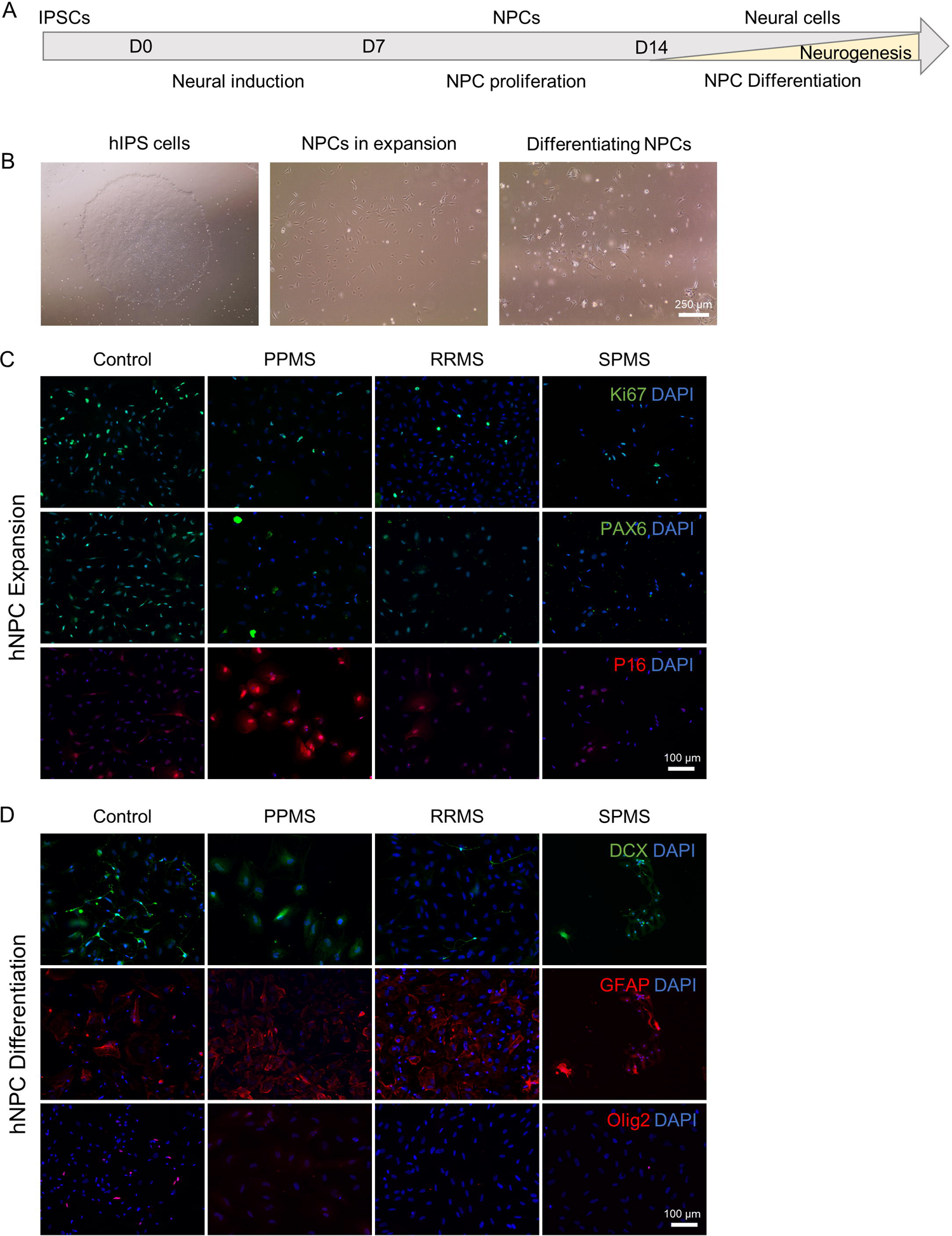
IPS derived NPCs reveal senescence and disrupted stem cell capacity. To confirm our findings about the neural precursor pool in c-organoids, patient derived iPSCs were directly differentiated into neural stem cells. Obtained patient NSCs were studied by immunofluorescence during expansion and after differentiation by mitogen withdrawal. A) Schematic representation of the protocol used to differentiate iPSCs into NPC. B) Representative pictures of iPSCs in expansion (left), NPCs in expansion (middle), and differentiated NPCs after 10 days of mitogen withdrawal (right). C) Representative pictures of immunofluorescence in NPCs during expansion in vitro. Staining for proliferation marker Ki67, stem cell marker PAX6 and senescence marker P16. A decrease of Ki67 and PAX6 expression was observed in MS organoids compared to control. A strong expression of P16 was detected in PPMS NPCs only. D) Representative pictures of immunofluorescence in NPCs after differentiation by mitogen withdrawal. Staining for neuronal marker DCX, astroglial marker GFAP and the oligodendroglial lineage marker Olig2 were performed. A decrease of DCX and Olig2, associated with an increase of GFAP expression was detected in MS samples, particularly PPMS.

First, an immunofluorescence against pluripotency marker Oct4 and Nanog was performed to verify that IPSC has lost pluripotency capacity and entered differentiation (data not shown). As expected only a very few cells were positive for those markers.

During expansion phase of IPS-derived NPCs, immunofluorescence was performed for proliferation marker Ki67 and stem cell marker PAX6 (Fig. 8C). A reduction of each marker was detected in MS samples, most notably in PPMS. In parallel, a staining for the senescence marker p16INK4A was performed. In control, RRMS and SPMS samples no/very low number of p16INK4A^+^ cells were detected while majority of cells in PPMS sample were expression this marker. This increase of senescence in PPMS NPCs compared to control and RRMS NPCs has previously been observed (Nicaise et al., 2019; Mutukula et al., 2021), highlighting cellular senescence in PPMS NPCs.

After 10 days of differentiation by mitogen withdrawal, immunofluorescence was performed for differentiation markers DCX, GFAP and Olig2 to detect difference in neuronal, astroglia and oligodendrocyte differentiation respectively (Fig. 8D). In control samples, some cells exhibited a neuronal morphology and were positive for neuronal marker DCX, but some cells were also positive for GFAP and Olig2 showing differentiation capacity in every neural cell type. In RRMS and SPMS NPCs, less cells seemed to express DCX or Olig2 while more cells expressed GFAP, and for PPMS NPCs, no DCX or Olig2 positive cell were found, the vast majority of the NPCs were expressing GFAP (Fig. 8D).

These results highlight a strong senescence of PPMS NPCs leading to a disruption of stemness of the progenitor pool, perhaps explaining the known diminished remyelination potential in PPMS (Nicaise et al., 2017; Nicaise et al., 2019).

## DISCUSSION

We describe c-organoids as an innovative model to study MS. We detected a decrease of proliferative capacity, notably in progressive forms of MS, associated with a reduction of the progenitor pool and an increase of neurogenesis possibly due to a symmetric shift of the mode cell division. We linked these effects to a strong decrease of p21 expression in PPMS organoids, unrelated to the DNA damage and apoptosis pathway.

Organoids derived from patient iPSCs retained the genetic information of the donor allowing analysis of the effect of genetic on the generation, proliferation and differentiation of neural progenitors into glial cells and neurons. It is important to note that c-organoids do not contain blood vessel or immune cells, and thus do not reproduce the inflammatory demyelination classically associated with MS. We used this as an advantage to understand the genetic variant liability in MS pathogenesis. Studies indicate that genetic and environmental factors interact to make certain individuals more susceptible to the disease. At present 1961 non-MHC variants in 156 genomic regions are significantly associated with MS and 21% of intergenic variants are associated with gene regulation in cortex tissue but not in immune cells (International Multiple Sclerosis Genetics, 2019) showing that c-organoids might be a very useful tool to study the effect of MS genetic variants on human corticogenesis.

For this study we derived IPSC from blood samples of patients with MS. The feasibility of creating IPSC lines from MS patients was reported with patient fibroblasts reprogrammed by retroviral delivery (Song et al., 2012; Miquel-Serra et al., 2017) or by mRNA/miRNA method (Douvaras et al., 2014) but also from blood cells using sendai virus (Nicaise et al., 2017; Mutukula et al., 2021) or from renal epithelial cells transfected with episomal factors (Massa et al., 2016). All techniques allow efficient reprogramming of iPSCs, which exhibited robust pluripotency associated with strong differentiation capacity. Moreover, iPSCs derived from MS patient can differentiate into mature astrocytes, oligodendrocytes and neurons with normal karyotypes (Song et al., 2012; Douvaras et al., 2014; Massa et al., 2016).

We chose to use iPSCs derived from patient blood samples (source Tisch MSRCNY, table S1) as it required a less painful and invasive procedure than fibroblast. Cell lines were reprogrammed using electroporation of episomal vector to produce transgene-free, virus-free human iPSCs. IPS cell line from the New York Stem Cell Foundation reprogramed from fibroblasts were also used for this project. In our hands, no significant difference was found between iPSCs derived from blood cells or fibroblast, which was also shown in previous report (Nicaise et al., 2019). All IPS cell line used in this study successfully passed quality control and karyotype analysis.

In our c-organoids, cyclin-dependent kinase inhibitor p21 was mostly expressed in the VZ which contains the stem cell pool. The loss of p21 expression was associated with a decrease of proliferation and a lower number of progenitor cells. Localization of p21 within the nucleus in both control and MS organoids, indicates that its function might be related to the maintenance of stem cell renewal, DNA repair or cell cycle arrest (Kreis et al., 2019). Furthermore, p21 is important for maintaining neural stem cell quiescence and self-renewal and its loss induces a quick exhaustion of NSC after a few passages in vitro (Kippin et al., 2005). Indeed, p21^-/-^ NSCs from mice initially displayed increased expansion followed by reduced self-renewal in vitro, (Kippin et al., 2005) which might explain the fastest growth rate of PPMS organoids at D2 in vitro followed by a similar growth rate compared to other conditions observed after D9 (Fig. 3A-B). We also observed an exhaustion of PPMS IPS-derived NSC in vitro after 4-5 passages compared to control and other subtypes of MS highlighted by a positive immunostaining for p16 (Fig. 11C). Our data supports these known functions as we observed a decrease of stem cell population and stem cell proliferation capacity associated with a decrease with p21 expression in PPMS organoids.

In parallel of this reduction of stemness, we observed a significant increase of neurogenesis (Fig. 6). It has been reported that DCX and NeuN protein expressions were increased in p21^−/−^ mice, demonstrating increased neuron proliferation (Pechnick et al., 2008) and thus, neurogenesis, which correlates with our results To verify the hypothesis of an imbalance of the proliferation/differentiation ratio in PPMS we analyzed the stem cell division modes. We observed a shift of the division mode to asymmetry in MS organoids compared to control, which indicates a reduction of NPC proliferative capacity, in favor of neurogenesis. Interestingly, p21 downregulation is associated with an increased length of the G1 phase leading to transition from symmetric proliferative to asymmetric neurogenic division (Watanabe et al., 2015). This result indicates that p21 decrease in MS organoids may lead to the proliferation/differentiation imbalance through a modification of the cell cycle length.

We further characterized differentiation potential of NPCs in oligodendrocytes. In response to demyelination, oligodendrocyte precursor cells (OPCs) divide and differentiate to regenerate oligodendrocyte population. However, remyelination fails during the later stages of MS and the underlying mechanism are poorly understood (Levine et al., 2001). Our analysis showed a decrease of OPCs marker Olig2 in MS organoids compared to control.

p21 is associated with p53 activation and apoptosis following DNA damage leading to cell growth arrest (Marques-Torrejon et al., 2013) or alternatively p21 can be activated by DNA damage, inhibiting p53 and inducing symmetric self-renewing divisions (Insinga et al., 2013). We did not detect any change in γH2AX expression in PPMS organoids compared to control, suggesting that no DNA damage occurred in our model. Furthermore, no difference in p53 and CC3 protein levels was seen in PPMS organoids compared to control. Overall, the DNA damage and apoptosis pathway was not activated in our MS organoids, and might not be involved with the reduced proliferation and stem cell number observed in PPMS suggesting that difference of p21 expression might be due to a genetic/epigenetic alteration in our patient samples. This absence of link between NSC exhaustion and activation of apoptosis pathway in p21^-/-^ NSC has already been shown (Kippin et al., 2005). Interestingly, it has been reported that p21 is required for the differentiation of oligodendrocytes and that animal knockdown for p21 exhibited hypomyelinated brain (Zezula et al., 2001). This function of p21 on differentiation is independent of its ability to control exit from the cell cycle.

p21 is also described as an autoimmunity suppressor (Santiago-Raber et al., 2001). p21 can modify cell cycle progression, replicative senescence of immune cells but also hemopoietic stem cell quiescence, and apoptosis (Arias et al., 2007) probably through NF-κB activation and IFNγ production in T-cells (Daszkiewicz et al., 2015). Also, p21-deficient mice showed increased in vitro and in vivo T cell cycling and activation, leading to mild autoimmune manifestations (Santiago-Raber et al., 2001). Furthermore, p21^-/-^ mice developed a lethal autoimmune syndrome characterized by the production of autoantibodies with expansion of memory B and CD4^+^ T cells (Santiuste et al., 2010).

In IPS derived NSCs, we found a high expression of p16 in PPMS but not in RRMS, SPMS or control as seen in previous studies (Nicaise et al., 2019; Mutukula et al., 2021). PPMS iPSCs can be directly differentiated into OPC and mature oligodendrocytes (Douvaras et al., 2014). However, when iPSCs were first differentiated in NPCs, the PPMS NPCs did not provide neuroprotection against demyelination in cuprizone-fed mice and did not support OPC differentiation in vitro (Nicaise et al., 2017). These results indicate that, unlike RRMS and SPMS, PPMS is associated with exhaustion of the progenitor pool and/or senescence of the NPCs leading to remyelination failure. Intriguingly, remyelination failure is a characteristic clinical feature that distinguishes PPMS at disease onset from RRMS.

Despite advantages of the use of organoids to study MS, some challenges remain. Variability of organoid development is a known confounding factor. However, recent improved protocols lead to very consistent organoid development (Sivitilli et al., 2020), with a variability that is close to human brain variability. New protocols have also been developed to create vascularized organoids (Mansour et al., 2018; Shi et al., 2020) or to include mature myelinating oligodendrocytes (Shaker et al., 2021). C-organoids mimics the development of the human brain while MS occurs around 20-40 years old. However, organoid represent a very useful tool to study MS pathogenesis before the onset of the disease. It has also been used to model other neurodegenerative disorders such as Alzheimer’s disease (Raja et al., 2016) or Parkinson’s disease (Kim et al., 2019). Also, control of the microenvironment provides an opportunity to study the interplay of a genetic background and specific effects of putative environmental triggers of MS such as infection with Epstein-Barr virus.

In conclusion this work is a proof of principle, showing the c-organoids derived from patients with MS can be used as an innovative tool to better understand the genetic basis for phenotypic differences seen in MS. Using this model, we identified p21 as a new protein of interest in the pathogenesis of progressive MS.

## MATERIALS AND METHODS

### Patient selection

Peripheral blood mononuclear cells (PBMCs) samples were obtained from clinically definite MS patients diagnosed according to the 2017 McDonald Criteria and a healthy control subject. All MS patients underwent neurological examinations and MRI imaging and were classified as having RRMS, SPMS or PPMS by board certified neurologists specializing in MS care. All protocols were approved by IRB and all donors provided their written informed consent for participation. Human IPS cells were generated from PBMCs of blood samples of donors. Other cell lines were also received from the New York Stem Cell Foundation. Donor information can be found in table S1.

### Reprogramming CD34+ progenitor cells

Human PBMC’s were isolated following the “STEMCELL Integrated Workflow for the Isolation, Expansion, and Reprogramming of CD34+ Progenitor Cells”. Briefly, blood samples were collected in Heparin-vacutainer tubes from donors ranging from 8-20 mL samples. CD34+ hematopoietic stem and progenitor cells were isolated from peripheral blood using the EasySep RosetteSep kit (STEMCELL) and expanded in vitro in CD34+ expansion media made of StemSpan SFEM II and CD34+ expansion supplements (STEMCELL) (Fig. S1A). After 7-10 days of culture, 1×106 cells were collected for reprogramming by electroporation using the Epi5 Episomal iPSC Reprogramming Kit (ThermoFisher) and Human CD34+ Cell Nucleofector Kit (Lonza) using a Nucleofector 2b Device (Lonza). After electroporation cells were cultured on Matrigel coated 6-well plates (100 µg/ml, Corning) in CD34+ expansion media. After 3 days, ReproTeSR (STEMCELL) was added to culture media for 2 more days. At day 7 cell were cultured in ReproTeSR media only. Media was changed on a daily basis. After 2-3 weeks, IPS cell colonies were isolated manually (Fig. S1B) and transferred to Matrigel coated 6-well plates containing mTeSR plus media (STEMCELL). Three subclones were created for each IPS cell line.

### Human induced pluripotent stem cells

Human iPSCs were cultured as previously described (Lancaster and Knoblich, 2014; Daviaud et al., 2018; Daviaud et al., 2019). iPSCs were plated in 6-well tissue culture plates coated with diluted Matrigel (100 µg/ml, Corning) and maintained in mTeSR Plus Culture Media (STEMCELL), supplemented with rock inhibitor Thiazovivin (2 µM, Millipore). Media was then changed on a daily basis without rock inhibitor until ready to passage at about 70-80% confluency or harvest.

All human pluripotent stem cells were maintained below passage 30 and confirmed negative for mycoplasma using the MycoFluor Mycoplasma Detection Kit (ThermoFisher). iPSCs were regularly tested for pluripotency using OCT4, NANOG and SOX2 markers (Fig. S1C) and were confirmed to be karyotypically normal by G-band testing (Fig. S1D).

### Generation of neural precursor cells

Neural precursor cells (NPCs) were produced following a protocol previously described (Gunhanlar et al., 2018) with minor modification. Briefly, iPSCs were dissociated with EDTA (0.5 mM, Millipore) for 5-6 minutes at 37°C. Embryonic Bodies (EBs) were generated by transferring 4000 cells per well of an ultra-low attachment 96 well plates in mTeSR Plus supplement with Thiazovivin (2 µM, Millipore). After 2 days, media was changed for neural induction media (StemDiff, STEMCELL) and EBs were cultured for 4 more days. At day 7, EBs were slightly dissociated by mechanical trituration and were put in culture on Matrigel (100 µg/ml, Corning) coated plates in neural induction media (StemDiff, STEMCELL) for 7 days. At d15 media was switched to NPC medium composed of DMEM/F12, 1% N2 supplement (ThermoFisher), 2% B27 with no RA supplement (ThermoFisher), 20 ng/ml epithelial growth factor (Peprotech), 10 ng/ml basic fibroblast growth factor (Peprotech) and 1% penicillin/streptomycin (ThermoFisher). On d15, cells were considered pre-NPCs and able to be passaged and cryopreserved when confluent. From passage 3, cells were considered NPCs and used for histological analysis and for neural differentiation.

Neural differentiation was achieved by mitogen withdrawal. NPCs were cultured for 10 days on Matrigel coated plates with differentiation media made of DMEM/F12, 1% N2 supplement and 2% B27 with RA supplement (ThermoFisher) and were then fixed and analyzed by immunostaining.

### Generation of cerebral organoids

C-organoids were generated from human iPSCs and processed for analysis as described (Lancaster and Knoblich, 2014; Daviaud et al., 2018; Daviaud et al., 2019) with minor modifications (Fig. S2A). iPSCs were washed with Dulbecco’s phosphate-buffered saline (ThermoFisher) and dissociated with EDTA 0.5 mM (Millipore) to generate single cells. A total of 4.5 x 103 cells were seeded into each well of an ultra-low-attachment 96-well plate (Corning) to form embryoid bodies in mTeSR Plus medium supplemented with 4µM of Thiazovivin (Millipore) for the first 2 days. Media was changed every other day to the same medium without Thiazovivin for another 2-3 days. After 4-5 days of culture or when embryonic bodies (EBs) reached ∼500-600 µm in diameter and when surface tissue began to brighten, EBs were cultured in neural induction medium (StemDiff, STEMCELL). After neuroepithelium emerged (usually at ∼ day 9-10), embryoid bodies were embedded in 15 µL Matrigel droplets and cultured in 6 well plates containing c-organoid differentiation media consisting of 1:1 DMEM-F12 and Neurobasal media (Gibco), with addition of 0.5% N2 supplement (Life Technologies), 0.5% ml MEM-NEAA (Gibco), 1% Glutamax (Gibco), 1% B27 supplement without Vitamin A (Life Technologies), 0.1 µM of 2-Mercaptoethanol (Millipore), 2.6 µg/ml Insulin (Sigma Aldrich) in static culture for 4 days. Organoids were then cultured in c-organoid differentiation media supplemented with Vitamin A on an orbital shaker (CO2 Resistant Shakers, ThermoFisher) at 80 rpm. Organoids were cultured for 42 days while observing radial growth and neuroepithelial bud formation.

### Histological Analysis

At D28 and D42 of culture, c-organoids were washed in D-PBS before fixing in 4% PFA (ThermoFisher) for 15 minutes at 4°C. After three washes with D-PBS, the c-organoids were cryoprotected in 30% sucrose overnight at 4°C followed by flash freezing in OCT Compound (ThermoFisher) and stored at −20°C. Cryosections of organoids were obtained at 15 µm thickness using a cryostat (Leica CM 1950) and mounted on microscope slides (Histobond+, VWR).

For immunofluorescence, slides were thawed to room temperature before being outlined by a PAP pen (Millipore) to create a hydrophobic barrier. Slides were washed and permeabilized with PBS supplemented with 0.1% Triton X-100 (Millipore Sigma). Non-specific binding sites were blocked with PBS supplemented with 0.1% Tween 20, 4% Bovine Serum Albumin (ThermoFisher) and 10% Normal Goat Serum (ThermoFisher) for 1 hour at RT. Slides were then incubated overnight at 4°C with the following primary antibodies diluted in blocking solution: rabbit anti-cleaved caspase 3 (1:200, Novus Biologicals), rat anti-CTIP2 (1:500, Abcam), guinea Pig anti-DCX (1:500, Millipore), mouse anti-EOMES (1:100, ThermoFisher), mouse anti-GAD67 (1:100, ThermoFisher), mouse anti-Ki67 (1:400, Millipore), mouse anti-Nanog (1:400, Abcam), rabbit anti-Olig2 (1:200, Abcam), rabbit anti-Oct4 (1:400, Abcam), rabbit anti-p16 (1:200, ThermoFisher), rabbit anti-p21 (1:200, ThermoFisher), rabbit anti-p53 (1:100, ThermoFisher), mouse anti-PAX6 (1:100, Abcam), rabbit anti-phospho-Histone H2AX (1:200, ThermoFisher), mouse anti-phospho-Histone H3 (1:400, Abcam), mouse anti-SOX2 (1:400, Abcam), rabbit anti-TBR1 (1:400, Abcam), rabbit anti-vGluT1 (1:200, Abcam). After washes, slides were incubated with appropriate Alexa-coupled secondary antibodies (ThermoFisher) diluted in blocking solution for 1 hour at RT and counterstained with DAPI, before mounting with Fluoromount Aqueous Mounting Medium (Millipore).

Immunofluorescence images were collected on a fluorescent microscope (Zeiss Imager M2) or a confocal fluorescent microscope (Zeiss LSM 510) and processed with Zen software and Fiji software (Schindelin et al., 2012).

### Experimental design and statistical analysis

C-organoids were derived from, at least, two different patient cell lines. For each cell line, 5-6 independent batches were made with 3-4 organoids per batch. Therefore, a total of 30-40 organoids were analyzed for this work and 1-3 representative images from each organoid were quantified (Fiji software) (Schindelin et al., 2012).

To determine the percentage of stained cells in the c-organoid sections, the appropriate immunohistochemistry markers counterstained with DAPI were used for quantification. For each section, regions of interest were generated in 250 µm (width) x 300 µm (height) radial columns spanning all cortical layers near organoid surface to normalize area of analysis.

Statistical analysis was performed using GraphPad prism 8. Differences between multiple conditions were determined using a one-way analysis of variance (ANOVA) test. If the ANOVA residuals were normally distributed (Shapiro-Wilk test), a Tukey follow-up test was then performed for multiple comparison. In the case that the ANOVA residuals were not normally distributed (Shapiro-Wilk test), a Kruskal-Wallis test was performed followed by a Dunn’s test for multiple comparison. Differences between grouped conditions were assessed using a Two-way ANOVA test, followed by a Tukey post hoc test for condition comparison. To analyze frequencies distribution a Chi-square test was performed. All data were presented as the mean value with standard error of the mean (SEM) unless otherwise stated. Results were considered significant when p < 0.05.

## ACKNOWLEDGMENTS

The authors thank the members of Tisch MSRCNY for helpful discussions, and the New York Stem Cell Foundation for providing patient with MS derived IPS cell lines.

## SUPPLEMENTARY INFORMATION

**Figure S1:** CD34+ derived IPS cells characterization and quality control. A) Brightfield picture of patient CD34+ cells in expansion. B) Representative images of IPS cell colonies growing 20 days after reprogramming. iPSCs colonies were manually scraped and replated. The yellow dotted line shows the distinction between iPSCs and cells that failed to reprogram. C) Characterization of iPSCs pluripotency markers. Left panel shows brightfield picture of IPS cells. Right panels show immunofluorescence for pluripotency markers Oct4, Nanog and the DAPI counterstain. D) Karyotype analysis of CD34+ derived iPSCs, all display normal karyotype.

**Figure S2:** C-organoids protocol and cell population characterization. A) Schematic representation (top panels) and brightfield pictures (bottom panels) of the main steps of the c-organoid derivation protocol. B) Immunofluorescence images of c-organoids at D42 demonstrate ventricle-like structure (V) aligned with neural precursors and their progeny at different stages of differentiation. Ki67 marker represents proliferating cells, SOX2 for neural stem cells, TBR2 for intermediate progenitors, DCX for neuroblasts, TBR1 for early-born cortical neurons and CTIP2 for deep-layer subcortical neurons. DAPI for nuclear counterstaining. C) Schematic representation of the cortical structure found in human c-organoids. The ventricular zone containing the stem cell pool and most proliferating cells, the SVZ containing mostly IPCs and the oSVZ and CP containing neuroblasts and mature neurons.

**Figure S3:** Glutamatergic and GABAergic neurons expression. A) Pictures of immunofluorescence against GABAergic (GAD67) and Glutamatergic (vGluT1) neuronal markers taken by confocal microscopy in control (top) and PPMS (bottom) in c-organoids at D42. The right panels show higher magnifications. White arrows show positive cells.

## REFERENCE

1. Andersson PB, Waubant E, Gee L, Goodkin DE (1999) Multiple sclerosis that is progressive from the time of onset: clinical characteristics and progression of disability. Arch Neurol 56:1138–1142.

2. Arias CF, Ballesteros-Tato A, Garcia MI, Martin-Caballero J, Flores JM, Martinez AC, Balomenos D (2007) p21CIP1/WAF1 controls proliferation of activated/memory T cells and affects homeostasis and memory T cell responses. J Immunol 178:2296–2306.

3. Cazzalini O, Scovassi AI, Savio M, Stivala LA, Prosperi E (2010) Multiple roles of the cell cycle inhibitor p21(CDKN1A) in the DNA damage response. Mutat Res 704:12–20.

4. Chenn A, McConnell SK (1995) Cleavage orientation and the asymmetric inheritance of Notch1 immunoreactivity in mammalian neurogenesis. Cell 82:631–641.

5. Daszkiewicz L, Vazquez-Mateo C, Rackov G, Ballesteros-Tato A, Weber K, Madrigal-Aviles A, Di Pilato M, Fotedar A, Fotedar R, Flores JM, Esteban M, Martinez AC, Balomenos D (2015) Distinct p21 requirements for regulating normal and self-reactive T cells through IFN-gamma production. Sci Rep 5:7691.

6. Daviaud N, Friedel RH, Zou H (2018) Vascularization and Engraftment of Transplanted Human Cerebral Organoids in Mouse Cortex. eNeuro 5.

7. Daviaud N, Chevalier C, Friedel RH, Zou H (2019) Distinct Vulnerability and Resilience of Human Neuroprogenitor Subtypes in Cerebral Organoid Model of Prenatal Hypoxic Injury. Front Cell Neurosci 13:336.

8. Douvaras P, Wang J, Zimmer M, Hanchuk S, O’Bara MA, Sadiq S, Sim FJ, Goldman J, Fossati V (2014) Efficient generation of myelinating oligodendrocytes from primary progressive multiple sclerosis patients by induced pluripotent stem cells. Stem Cell Reports 3:250–259.

9. Englund C, Fink A, Lau C, Pham D, Daza RA, Bulfone A, Kowalczyk T, Hevner RF (2005) Pax6, Tbr2, and Tbr1 are expressed sequentially by radial glia, intermediate progenitor cells, and postmitotic neurons in developing neocortex. J Neurosci 25:247-251.

10. Filippi M, Bar-Or A, Piehl F, Preziosa P, Solari A, Vukusic S, Rocca MA (2018) Multiple sclerosis. Nat Rev Dis Primers 4:43.

11. Gleeson JG, Lin PT, Flanagan LA, Walsh CA (1999) Doublecortin is a microtubule-associated protein and is expressed widely by migrating neurons. Neuron 23:257–271.

12. Hodge RD, Kahoud RJ, Hevner RF (2012) Transcriptional control of glutamatergic differentiation during adult neurogenesis. Cell Mol Life Sci 69:2125–2134.

13. Insinga A, Cicalese A, Faretta M, Gallo B, Albano L, Ronzoni S, Furia L, Viale A, Pelicci PG (2013) DNA damage in stem cells activates p21, inhibits p53, and induces symmetric self-renewing divisions. Proc Natl Acad Sci U S A 110:3931–3936.

14. International Multiple Sclerosis Genetics C (2019) Multiple sclerosis genomic map implicates peripheral immune cells and microglia in susceptibility. Science 365.

15. Kim H, Park HJ, Choi H, Chang Y, Park H, Shin J, Kim J, Lengner CJ, Lee YK, Kim J (2019) Modeling G2019S-LRRK2 Sporadic Parkinson’s Disease in 3D Midbrain Organoids. Stem Cell Reports 12:518–531.

16. Kippin TE, Martens DJ, van der Kooy D (2005) p21 loss compromises the relative quiescence of forebrain stem cell proliferation leading to exhaustion of their proliferation capacity. Genes Dev 19:756–767.

17. Kreis NN, Louwen F, Yuan J (2019) The Multifaceted p21 (Cip1/Waf1/CDKN1A) in Cell Differentiation, Migration and Cancer Therapy. Cancers (Basel) 11.

18. Lancaster MA, Knoblich JA (2014) Generation of cerebral organoids from human pluripotent stem cells. Nat Protoc 9:2329–2340.

19. Lancaster MA, Renner M, Martin CA, Wenzel D, Bicknell LS, Hurles ME, Homfray T, Penninger JM, Jackson AP, Knoblich JA (2013) Cerebral organoids model human brain development and microcephaly. Nature 501:373–379.

20. Levine JM, Reynolds R, Fawcett JW (2001) The oligodendrocyte precursor cell in health and disease. Trends Neurosci 24:39–47.

21. Lublin FD, Reingold SC (1996) Defining the clinical course of multiple sclerosis: results of an international survey. National Multiple Sclerosis Society (USA) Advisory Committee on Clinical Trials of New Agents in Multiple Sclerosis. Neurology 46:907–911.

22. Macleod KF, Sherry N, Hannon G, Beach D, Tokino T, Kinzler K, Vogelstein B, Jacks T (1995) p53-dependent and independent expression of p21 during cell growth, differentiation, and DNA damage. Genes Dev 9:935–944.

23. Madhavan M, Nevin ZS, Shick HE, Garrison E, Clarkson-Paredes C, Karl M, Clayton BLL, Factor DC, Allan KC, Barbar L, Jain T, Douvaras P, Fossati V, Miller RH, Tesar PJ (2018) Induction of myelinating oligodendrocytes in human cortical spheroids. Nat Methods 15:700–706.

24. Mansour AA, Goncalves JT, Bloyd CW, Li H, Fernandes S, Quang D, Johnston S, Parylak SL, Jin X, Gage FH (2018) An in vivo model of functional and vascularized human brain organoids. Nat Biotechnol 36:432–441.

25. Marques-Torrejon MA, Porlan E, Banito A, Gomez-Ibarlucea E, Lopez-Contreras AJ, Fernandez-Capetillo O, Vidal A, Gil J, Torres J, Farinas I (2013) Cyclin-dependent kinase inhibitor p21 controls adult neural stem cell expansion by regulating Sox2 gene expression. Cell Stem Cell 12:88–100.

26. Massa MG, Gisevius B, Hirschberg S, Hinz L, Schmidt M, Gold R, Prochnow N, Haghikia A (2016) Multiple Sclerosis Patient-Specific Primary Neurons Differentiated from Urinary Renal Epithelial Cells via Induced Pluripotent Stem Cells. PLoS One 11:e0155274.

27. Matsui TK, Matsubayashi M, Sakaguchi YM, Hayashi RK, Zheng C, Sugie K, Hasegawa M, Nakagawa T, Mori E (2018) Six-month cultured cerebral organoids from human ES cells contain matured neural cells. Neurosci Lett 670:75–82.

28. Mione MC, Cavanagh JF, Harris B, Parnavelas JG (1997) Cell fate specification and symmetrical/asymmetrical divisions in the developing cerebral cortex. J Neurosci 17:2018–2029.

29. Miquel-Serra L, Duarri A, Munoz Y, Kuebler B, Aran B, Costa C, Marti M, Comabella M, Malhotra S, Montalban X, Veiga A, Raya A (2017) Generation of six multiple sclerosis patient-derived induced pluripotent stem cell lines. Stem Cell Res 24:155–159.

30. Mutukula N, Man Z, Takahashi Y, Iniesta Martinez F, Morales M, Carreon-Guarnizo E, Hernandez Clares R, Garcia-Bernal D, Martinez Martinez L, Lajara J, Nunez Delicado E, Meca Lallana JE, Izpisua Belmonte JC (2021) Generation of RRMS and PPMS specific iPSCs as a platform for modeling Multiple Sclerosis. Stem Cell Res 53:102319.

31. Nadadhur AG, Emperador Melero J, Meijer M, Schut D, Jacobs G, Li KW, Hjorth JJJ, Meredith RM, Toonen RF, Van Kesteren RE, Smit AB, Verhage M, Heine VM (2017) Multi-level characterization of balanced inhibitory-excitatory cortical neuron network derived from human pluripotent stem cells. PLoS One 12:e0178533.

32. Nicaise AM, Banda E, Guzzo RM, Russomanno K, Castro-Borrero W, Willis CM, Johnson KM, Lo AC, Crocker SJ (2017) iPS-derived neural progenitor cells from PPMS patients reveal defect in myelin injury response. Exp Neurol 288:114–121.

33. Nicaise AM, Wagstaff LJ, Willis CM, Paisie C, Chandok H, Robson P, Fossati V, Williams A, Crocker SJ (2019) Cellular senescence in progenitor cells contributes to diminished remyelination potential in progressive multiple sclerosis. Proc Natl Acad Sci U S A 116:9030–9039.

34. O’Gorman C, Lin R, Stankovich J, Broadley SA (2013) Modelling genetic susceptibility to multiple sclerosis with family data. Neuroepidemiology 40:1–12.

35. Pechnick RN, Zonis S, Wawrowsky K, Pourmorady J, Chesnokova V (2008) p21Cip1 restricts neuronal proliferation in the subgranular zone of the dentate gyrus of the hippocampus. Proc Natl Acad Sci U S A 105:1358–1363.

36. Raja WK, Mungenast AE, Lin YT, Ko T, Abdurrob F, Seo J, Tsai LH (2016) Self-Organizing 3D Human Neural Tissue Derived from Induced Pluripotent Stem Cells Recapitulate Alzheimer’s Disease Phenotypes. PLoS One 11:e0161969.

37. Ransohoff RM (2012) Animal models of multiple sclerosis: the good, the bad and the bottom line. Nat Neurosci 15:1074–1077.

38. Sadovnick AD, Yee IM, Ebers GC, Risch NJ (1998) Effect of age at onset and parental disease status on sibling risks for MS. Neurology 50:719–723.

39. Santiago-Raber ML, Lawson BR, Dummer W, Barnhouse M, Koundouris S, Wilson CB, Kono DH, Theofilopoulos AN (2001) Role of cyclin kinase inhibitor p21 in systemic autoimmunity. J Immunol 167:4067–4074.

40. Santiuste I, Buelta L, Iglesias M, Genre F, Mazorra F, Izui S, Merino J, Merino R (2010) B-cell overexpression of Bcl-2 cooperates with p21 deficiency for the induction of autoimmunity and lymphomas. J Autoimmun 35:316–324.

41. Sessa A, Mao CA, Hadjantonakis AK, Klein WH, Broccoli V (2008) Tbr2 directs conversion of radial glia into basal precursors and guides neuronal amplification by indirect neurogenesis in the developing neocortex. Neuron 60:56–69.

42. Shaker MR, Pietrogrande G, Martin S, Lee JH, Sun W, Wolvetang EJ (2021) Rapid and Efficient Generation of Myelinating Human Oligodendrocytes in Organoids. Front Cell Neurosci 15:631548.

43. Shi Y, Sun L, Wang M, Liu J, Zhong S, Li R, Li P, Guo L, Fang A, Chen R, Ge WP, Wu Q, Wang X (2020) Vascularized human cortical organoids (vOrganoids) model cortical development in vivo. PLoS Biol 18:e3000705.

44. Sivitilli AA, Gosio JT, Ghoshal B, Evstratova A, Trcka D, Ghiasi P, Hernandez JJ, Beaulieu JM, Wrana JL, Attisano L (2020) Robust production of uniform human cerebral organoids from pluripotent stem cells. Life Sci Alliance 3.

45. Song B, Sun G, Herszfeld D, Sylvain A, Campanale NV, Hirst CE, Caine S, Parkington HC, Tonta MA, Coleman HA, Short M, Ricardo SD, Reubinoff B, Bernard CC (2012) Neural differentiation of patient specific iPS cells as a novel approach to study the pathophysiology of multiple sclerosis. Stem Cell Res 8:259–273.

46. Watanabe N, Kageyama R, Ohtsuka T (2015) Hbp1 regulates the timing of neuronal differentiation during cortical development by controlling cell cycle progression. Development 142:2278–2290.

47. Westerlind H, Bostrom I, Stawiarz L, Landtblom AM, Almqvist C, Hillert J (2014) New data identify an increasing sex ratio of multiple sclerosis in Sweden. Mult Scler 20:1578–1583.

48. Willer CJ, Dyment DA, Risch NJ, Sadovnick AD, Ebers GC, Canadian Collaborative Study G (2003) Twin concordance and sibling recurrence rates in multiple sclerosis. Proc Natl Acad Sci U S A 100:12877–12882.

49. Zezula J, Casaccia-Bonnefil P, Ezhevsky SA, Osterhout DJ, Levine JM, Dowdy SF, Chao MV, Koff A (2001) p21cip1 is required for the differentiation of oligodendrocytes independently of cell cycle withdrawal. EMBO Rep 2:27–34.

